# Face-like holistic Processing in Non-Face Stimuli

**DOI:** 10.1101/2025.03.07.641889

**Authors:** Yuxuan Zeng, Zitong Lu, Ren Hentz, David Osher

## Abstract

Holistic processing of visual stimuli has long been regarded as unique to faces, or otherwise extended to other object categories given sufficient expertise. We designed novel abstract stimuli that are recognizable strictly by configural information, and matched control stimuli that require the use of only featural information. We then tested four classic markers of holistic perception: inversion, misalignment, part–whole, and composite effects. We found that second-order configuration stimuli elicited robust holistic effects, while featural stimuli do not, showing that such effects emerge specifically when recognition depends on configural information, but not when it relies on featural cues. We further observed that first-order configuration stimuli were also sufficient to induce holistic effects, indicating that holistic processing can emerge from multiple levels of configural information. We also found a significant correlation between individual differences in face recognition ability and holistic processing effects with the second-order configuration stimuli, but not with the first order configuration stimuli, nor the featural stimuli. Lastly, convolutional neural networks trained on the same stimuli reproduced these patterns, strengthening the interpretation that holistic processing arises when configural information must be used to recognize a stimulus. Together, these findings demonstrate that holistic processing is fundamentally rooted in the representation of spatial configuration, with second-order relations providing the critical link to face recognition ability.

## 1. Introduction

Humans perceive faces in a holistic way, meaning we tend to recognize the face as a unified whole, rather than as a collection of individual features (Galton, 1879). This is often contrasted with how we perceive other objects, and holistic processing has long been considered a behavioral hallmark of face perception (Jiang et al., 2011; Yovel & Kanwisher, 2004, 2005).

Since Tanaka and Farah’s (1993) influential proposal that faces are recognized as templates, a wide range of behavioral effects, such as the inversion effect, part-whole effect, and composite effect, have shown that faces are processed as the configuration of features rather than processing each feature separately from the whole. Hence, holistic processing for faces is sometimes used interchangeably with the term ‘configural processing’ and these inversion, part-whole, and composite effects are essentially the behavioral markers of face processing. However, these markers have not been consistently tested for non-face stimulus classes or for novel inanimate stimulus classes that cannot rely on specialized processing mechanisms for faces or bodies, leaving open the question of whether holistic processing can be evoked for other stimulus classes. Here, we comprehensively test the behavioral hallmarks of face perception on a novel set of stimuli that can be discriminated only via configural properties and compare these effects to virtually identical control stimuli that can be discriminated only via featural properties. By generating these novel stimuli, we can test both first and second order configural effects on face-like holistic processing on non-face or non-animate stimuli that do not evoke a particular categorical representation (e.g. houses, chairs). In addition, we examine whether first- and second-order configural holistic processing is related to individual differences in face-recognition ability, providing insight into the mechanisms on which face recognition relies when processing non-face stimuli. If we are able to generate a set of stimuli that elicit face-like holistic effects, we will be better able to break down different components of holistic processing to understand its mechanisms without triggering e.g. face templates. This will get us closer to offering a better mechanistic explanation of why faces are a special domain in natural visual processing.

We test four classic paradigms that have been widely used to measure holistic face processing: the inversion, part-whole, composite, and misalignment tasks (Fig. 1a). The inversion effect, in which recognition drops significantly when a face is turned upside-down, is one of the earliest and most robust demonstrations of holistic disruption (Yin, 1969). Some researchers interpret this as a loss of holistic efficiency (Curby & Gauthier, 2009), while others argue inversion fundamentally breaks down configural integration (Rossion & Boremanse, 2008; Young et al., 1987). The part-whole effect shows that facial features (e.g., the nose) are better recognized when presented within the context of the whole face than in isolation, suggesting that faces are processed as integrated wholes (Tanaka & Farah, 1993). The composite effect goes a step further: when the top half of a face is combined with a mismatched bottom half, recognition is impaired because the two halves are automatically fused into a single holistic percept (Young et al., 1987).

**Figure 1.**
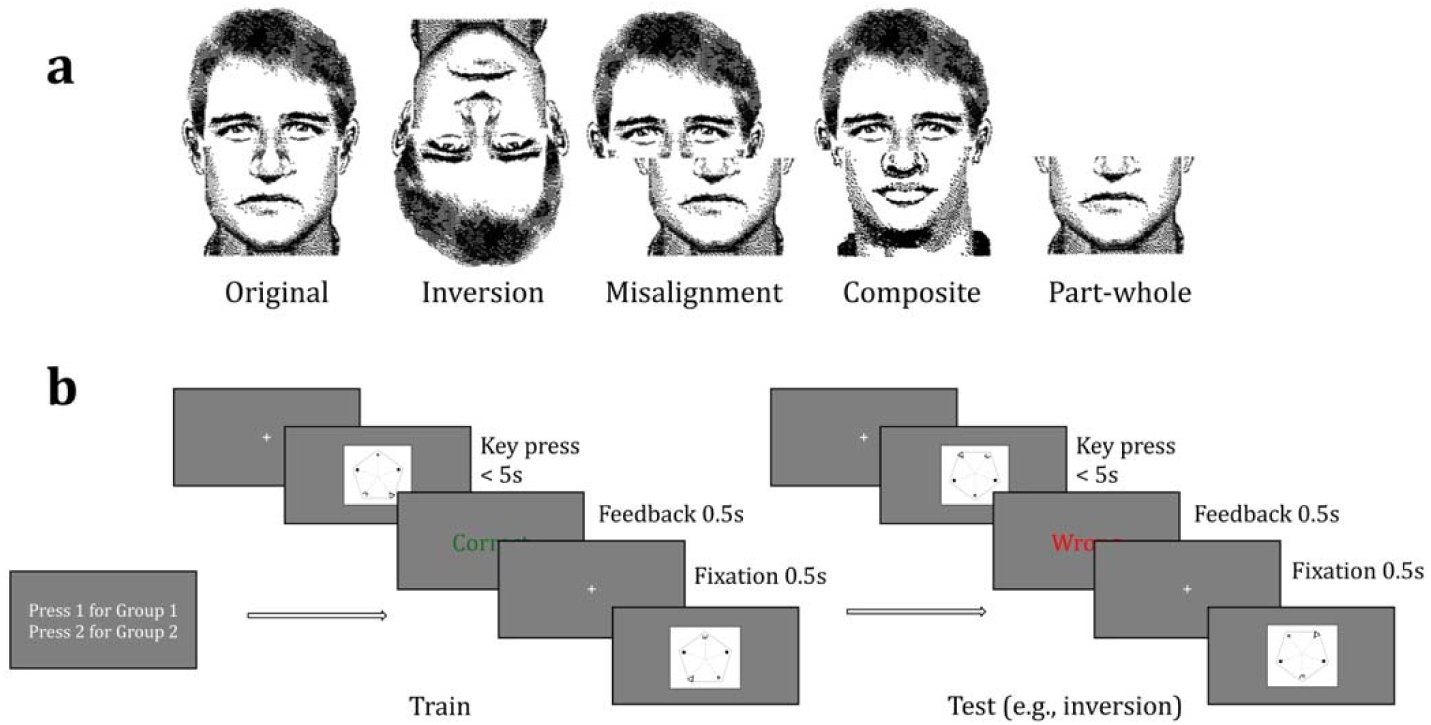
Experimental paradigm. (a) Several effects associated with holistic face processing, illustrated with faces: original, inversion, misalignment, composite, and part–whole (faces adapted from Farah et al., 1998; Farah, Tanaka, & Drain, 1995). (b) Behavioral paradigm. Participants first completed 300 training trials, followed by four tasks. Before each task began, participants were informed about the specific manipulation (e.g., stimuli would be inverted).

Misalignment itself also disrupts holistic processing and can thus serve as an independent probe. Each of these tasks captures a different facet of holistic processing, and prior work suggests they may reflect partially dissociable mechanisms (Rezlescu et al., 2017; Richler & Gauthier, 2014). To rigorously establish whether our stimuli elicited holistic processing, we systematically examined all hallmark paradigms associated with the construct. If consistent effects emerge across these distinct tasks, such convergence would offer strong evidence that our stimuli genuinely engaged holistic mechanisms. In addition, we examined whether performance on these four paradigms correlates with face-recognition ability as measured by the CFMT (Duchaine & Nakayama, 2006). Prior studies have reported mixed findings regarding whether holistic face-processing tasks reliably predict face-recognition performance (White & Burton, 2022), perhaps because both sides of the correlation involved face stimuli. If we find consistent associations between the four paradigms and face recognition in our non-face stimulus set, this would provide stronger evidence that holistic processing, independent of face-specific content, constitutes a foundational mechanism of face recognition.

In the present study, we designed novel, abstract stimuli that shared no physical resemblance to faces or known objects and could not be interpreted as animate objects (Fig 2a). These stimuli came in two versions: a configural condition, in which categorization depended solely on spatial relationships among features, and as a control, a featural condition, in which categorization was determined by which features were present, regardless of their spatial layout. If either of these non-face conditions elicits the same effects as observed for faces, it would challenge the notion that holistic processing is exclusive to face perception; and if the effects are larger for the configural condition it would suggest that the configural properties of stimuli are what drive the holistic processing effects we observe.

**Figure 2.**
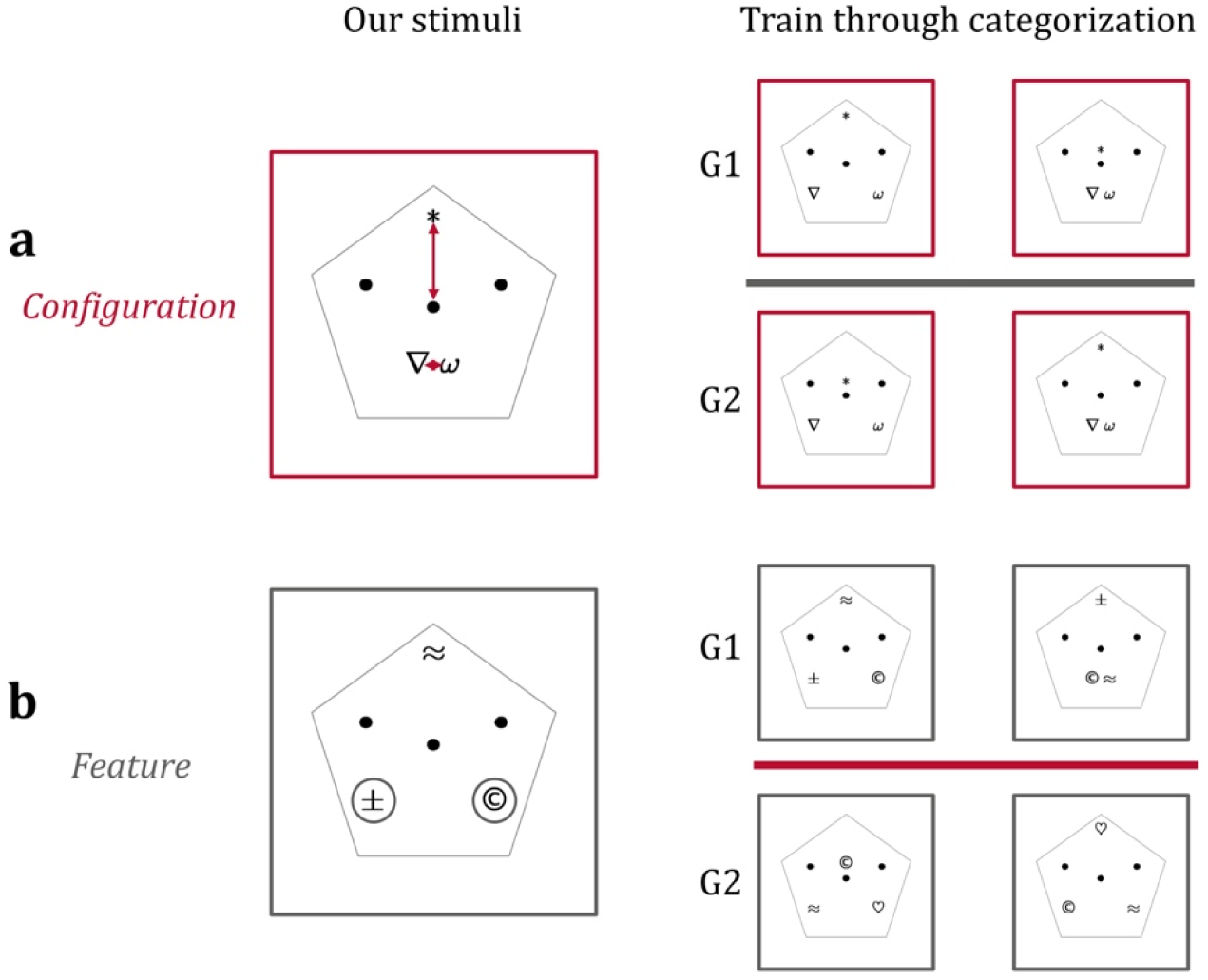
Examples of second-order stimuli. (cropped into squares for better visualization). (a) Configural stimuli. The red arrow illustrates the basis for grouping (not shown in the actual stimuli). Group 1 consisted of “long & wide” or “short & narrow” configurations, while Group 2 consisted of “short & wide” or “long & narrow” configurations. Because the grouping required the intersection of two configural dimensions, participants had to use configural information from both to categorize correctly. (b) Featural control stimuli. The gray circles illustrate the basis for grouping (not shown in the actual stimuli). Group 1 required the presence of both “©” and “±,” while Group 2 required the presence of both “♡” and “≈.” Since the third distractor symbol was drawn from the opposite group, correct categorization required recognizing at least two symbols.

We tested the hypothesis that second-order configural information (i.e., subtle spatial distances between features) is the key driver (Piepers & Robbins, 2012). Since all faces share the same first-order layout (e.g., eyes above nose, Tanaka & Sengco, 1997), second-order (e.g., the precise distance between the eyes) differences are critical for facial recognition (Diamond & Carey, 1986; McKone & Robbins, 2011), and some argue this is why faces uniquely engage holistic mechanisms. By generating non-face stimuli and manipulating first and second-order configurations, we are able to test this hypothesis without triggering the ‘face template’. If second-order configuration is necessary to trigger holistic processing, then only second-order stimuli should produce these effects.

An alternative hypothesis posits that holistic processing develops through extensive experience individuating objects within a category. According to this expertise account, faces elicit holistic perception not because of their configural properties per se, but because we have spent a lifetime learning to recognize them. Indeed, certain holistic effects have been observed for objects of expertise such as cars (Gauthier et al., 2000), dogs (Diamond & Carey, 1986), and even novel lab-trained animate stimuli like Greebles (Gauthier & Tarr, 1997). If expertise is a key driver of holistic processing, then minimal training (∼30 minutes in our case) on entirely novel stimuli with no resemblance to known object classes should be insufficient to induce holistic processing, for either the configural or featural stimulus sets.

Our results showed clear holistic effects for both first-order and second-order configuration conditions, but importantly, not for the featural control in either case. To further validate our findings, we trained a convolutional neural network (CNN) on the same tasks and found similar patterns of holistic behavior. These findings strongly suggest that holistic processing arises from configural information itself, rather than being exclusive to faces or requiring extensive expertise. As a further step, we also examined individual differences and found that only second-order configuration was reliably associated with face-recognition ability, a pattern consistent with classic findings. Together, these results provide a comprehensive account of how configural information underlies holistic processing and clarify its role in making faces a special domain of visual expertise.

## 2. Experiment 1: 2^nd^ Order Configuration

### 2.1 Human Behavior

#### 2.1.1 Methods

##### 2.1.1.1 Participants

Thirty-two undergraduates from The Ohio State University (OSU) participated in this experiment. They were recruited through the Research Experience Program (REP) and received course credits for their participation. Only those achieving an 80% accuracy rate in the last 30 trials of the preliminary training were included in the analysis, leaving 20 participants in the final sample.

##### 2.1.1.2 Stimuli

Stimuli were created using MATLAB and displayed on a standard 23.8-inch Dell monitor with a resolution of 1920×1080 pixels. Each stimulus consisted of an enclosed pentagon with five symbols (1167×875 pixels). Two fixed symbols (“●”) served as placeholders for positional uniformity across conditions, while the remaining three symbols determined group classification. No symbols were shared between the configural and featural conditions to ensure distinct grouping criteria:

- **Configural Condition** (Fig. 2a): Each stimulus contained three fixed symbols (“∇”, “*”, “ω”) whose relative spatial configuration (1^st^ order configuration) remained constant across all stimuli: the “*” symbol was always positioned at the top of the pentagon, “∇” at the lower-left, and “ω” at the lower-right. Group classification was determined by two second-order configural metrics: (1) the distance between the “*” symbol and the geometric center of the pentagon (categorized as either *long* or *short*), and (2) the distance between the “∇” and “ω” symbols (categorized as either *wide* or *narrow*). Group 1 stimuli were defined by either both *long & wide* or *short & narrow* distances, while Group 2 stimuli were defined by the combinations *short & wide* or *long & narrow*. Thus, successful classification required integration of **two** spatial relationships; neither distance alone was diagnostic. This design ensured that participants learned group structure based specifically on second-order configural information.
- **Featural Condition** (Fig. 2b): Featural stimuli provide a matched control for the configural stimuli since they require no configural information for recognition, and participants categorize them based on the features present alone. The stimuli preserved the same symbol positions used in the second-order configuration condition but replaced the original symbols with three drawn from a distinct feature set: “©”, “±”, “♡”, and “≈”. The spatial layout of these symbols was not predictive of group membership. Instead, classification was based purely on symbol identity: stimuli containing **both** “©” and “±” were assigned to Group 1, while those containing **both** “♡” and “≈” were assigned to Group 2. The third symbol was a distractor randomly selected from the opposite group. Importantly, group membership could only be determined by detecting specific **two** features rather than the presence of a single diagnostic symbol.

##### 2.1.1.3 Design and Procedure

The study employed a within-subject design, with participants completing both the configural and featural condition tasks in a randomized order. Within each condition, participants first completed a training phase with 300 trials to learn the categorization rules through feedback learning. In each trial, participants were shown a single stimulus and were instructed to decide whether they belong to “group 1” or “group 2” based on trial and error, responding via keyboard press. Stimuli were presented on the screen for up to 5 seconds or until a key response was made. If they responded within the time limit, feedback (“correct” or “wrong”) appeared immediately; if no response was made, “no response” was displayed. A fixation cross (“+”) was shown for 0.5 seconds between trials (Fig. 1b).

After training, participants progressed to a testing phase that was identical in procedure to the training task, except that the stimuli were altered in four ways: inverted (rotated 180°), misaligned, composite, and part (Fig. 3b). Prior to each task, participants were informed about these upcoming changes (e.g., “you will now see only part of the original stimuli”) and instructed to continue applying the classification rules learned during training. The four task conditions were presented in a randomized order, with each condition consisting of a block of 30 trials.

**Figure 3.**
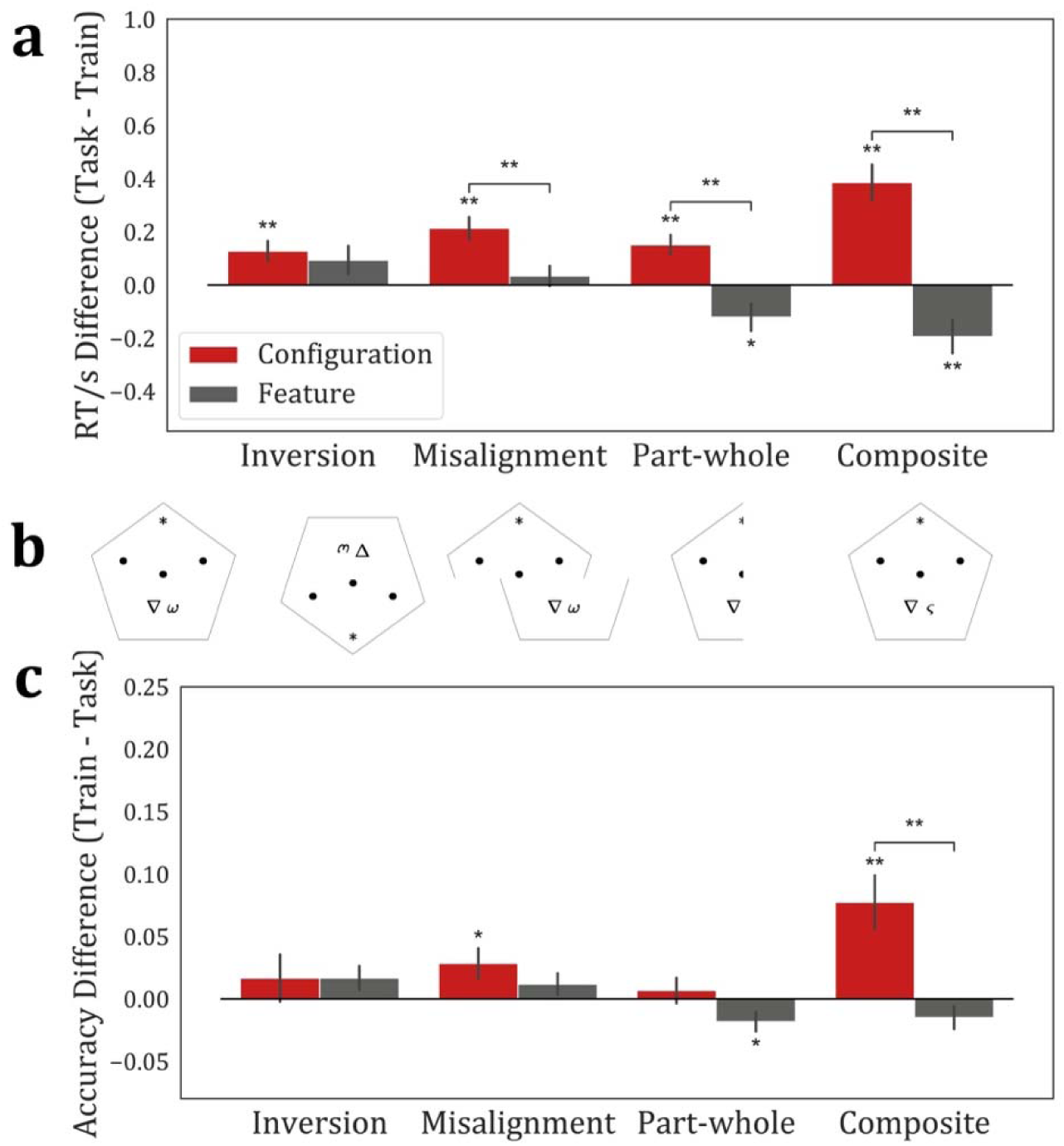
Behavioral results for second-order stimuli. (a) RT differences between training and task performance (Task RT – Train RT, plotted so that effects are primarily positive). (b) Example stimuli used during training and tasks. For the composite condition, the bottom-right symbol was replaced with a novel symbol, and its distance to the vertical centerline was randomized. (c) Accuracy differences between training and task performance (Train accuracy – Task accuracy, plotted so that effects are primarily positive). *, p < 0.05; **, significant after Bonferroni–Holm correction (p < 0.05).

Accuracy and reaction times from these trials were compared to the final 30 trials of the training phase to assess the presence and magnitude of holistic processing effects.

#### 2.1.2 Results

At the end of the training phase (last 30 trials), there were no significant differences between the configural and featural conditions for either RT or accuracy. RT was comparable across the two conditions, *t*(19) = −0.74, *p* = .468, and accuracy also did not differ significantly, *t*(19) = −1.88, *p* = .076. Mean RT was 0.98 s (SE = 0.06) in the configural condition and 1.03 s (SE = 0.07) in the featural condition. Mean accuracy was 95.50% (SE = 0.85%) for the configural condition and 97.17% (SE = 0.74%) for the featural condition. Therefore, the configural and featural conditions are most likely of similar difficulty and the featural condition is an adequate control for the configural condition.

In the testing phase, significant holistic processing effects emerged exclusively for the configural condition. RTs were significantly longer than baseline across all four tasks: inversion, t(19) = 3.38, p = .003; misalignment, t(19) = 5.03, p < .001; part–whole, t(19) = 4.00, p < .001; and composite, t(19) = 5.69, p < .001. In contrast, the featural control condition showed no significant effects for inversion, t(19) = 1.76, p = .094; misalignment, t(19) = 0.90, p = .380; or part–whole, t(19) = –2.39, p = .027 (not significant after Bonferroni–Holm correction).

Direct comparisons between the two conditions confirmed significant differences for the misalignment, part–whole, and composite tasks: misalignment, t(19) = 3.93, p < .001; part– whole, t(19) = 4.48, p < .001; and composite, t(19) = 5.82, p < .001. The inversion task showed no significant difference between conditions, t(19) = 0.61, p = .551 (Fig. 3a).

For accuracy, significant holistic effects emerged for the configural condition in the composite task, t(19) = 3.60, p = .002, indicating lower performance compared to the training baseline. The misalignment task, t(19) = 2.33, p = .031, did not survive Bonferroni–Holm correction but was significant as an independent test. The inversion, t(19) = 0.89, p = .387, and part–whole, t(19) = 0.66, p = .515, tasks showed no significant changes. In the featural control condition, no significant accuracy effects were observed after Bonferroni–Holm correction, including inversion, t(19) = 1.76, p = .096; misalignment, t(19) = 1.33, p = .199; part–whole, t(19) = –2.33, p = .031; and composite, t(19) = –1.63, p = .119.

Direct comparisons between the two conditions revealed a significant difference in the composite task, t(19) = 3.56, p = .002, while the other tasks (inversion, misalignment, and part–whole) did not differ reliably between conditions (all p > .05) (Fig. 3c).

### 2.2 Machine vision

In recent years, convolutional neural networks (CNNs) have achieved remarkable success in object recognition, which also raises questions about whether such models can capture processing mechanisms similar to those observed in human cognition. While CNNs are designed to excel in complex pattern recognition, it remains an open question whether they can replicate the same holistic effects that we observe in human visual perception tasks.

#### 2.2.1 Methods

##### 2.2.1.1 Stimuli

To generate a large number of images based on the original exemplars used in human behavior tasks to train and test CNN models, we implemented data augmentation by applying adding 5% random pixel noise. We generated 720 training images for each category (1440 images for configural task and 1440 images for featural task). Accordingly, we also generated corresponding upright (same data augmentation processing on the original exemplars but different from the images in the training set), inverted, misaligned, part, and composite images for the model test.

##### 2.2.1.2 Model and training procedure

We utilized pre-trained ResNet-18 models (He et al., 2015) on ImageNet (Russakovsky et al., 2015) to perform transfer learning based on our stimuli (for other ResNet models results, see supplemental material Figure S1), allowing them to learn the configuration and feature-based categorical classification tasks, resulting in a separate set of configuration-directed and feature-directed models. In the training of transfer learning, the models were trained using the standard categorical cross entropy loss with a static learning rate of 0.0002 and the Adam optimizer (Kingma & Ba, 2014). We trained 20 configuration-directed and 20 feature-directed ResNets to simulate 20 individuals’ behaviors.

##### 2.2.1.3 Model evaluations

We tested our configuration-directed and feature-directed ResNets on corresponding five types of test sets respectively. All ResNets achieved an accuracy of over 95% on the training stimuli. We computed the difference of classification accuracy between tasks and upright condition as a counterpart measure for human accuracy differences between tasks and training, and we used classification probability differences as a counterpart measure similar to human RT differences (Ratcliff & McKoon, 2008), for the inverted, misaligned, part, and composite conditions.

#### 2.2.2 Results

In the model testing phase, robust holistic processing effects emerged in the configural condition across all four tasks. Classification probabilities significantly deviated from the training baseline for configural networks in the inversion, *t*(19) = 16.44, *p* < .001; misalignment, *t*(19) = 20.19, *p* < .001; part–whole, *t*(19) = 12.32, *p* < .001; and composite, *t*(19) = 11.16, *p* < .001, conditions. In the featural condition, only inversion, *t*(19) = 11.16, *p* < .001, and misalignment, *t*(19) = 8.43, *p* < .001, produced significant deviations from baseline, but the magnitude of these changes was substantially smaller than in the configural condition. Both the part-whole task, *t*(19) = −3.85, *p* = .001, and composite task, *t*(19) = −68.60, *p* < .001, showed a strong reversal. Direct comparisons between conditions confirmed significantly larger effects for configural networks across all tasks: inversion, *t*(19) = 7.35, *p* < .001; misalignment, *t*(19) = 15.10, *p* < .001; part– whole, *t*(19) = 11.85, *p* < .001; and composite, *t*(19) = 63.17, *p* < .001 (Fig. 4a).

**Figure 4.**
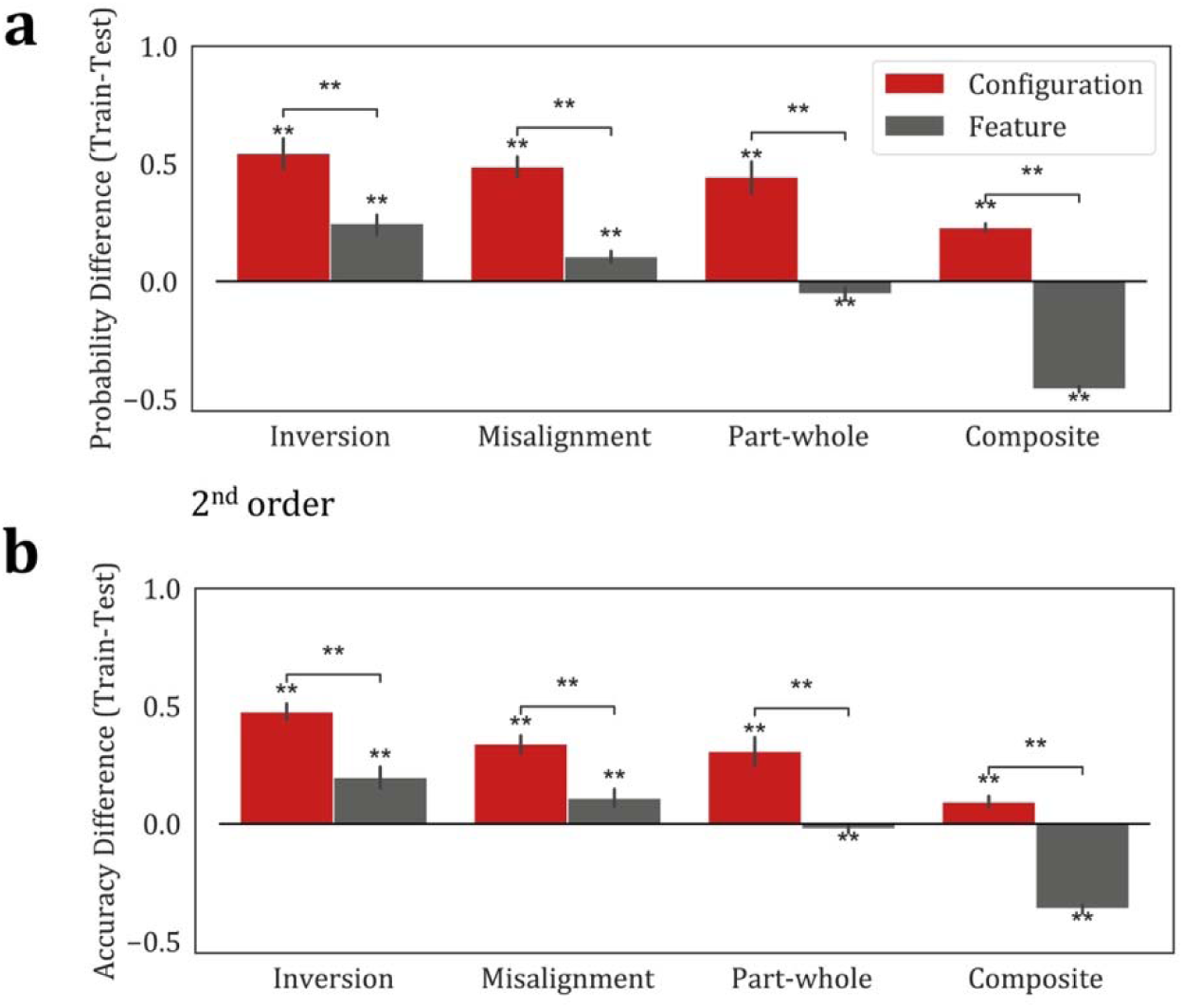
Computer vision results for second-order stimuli. (a) Probability differences between training and task performance (Train probability – Task probability, plotted so that effects are primarily positive). (b) Accuracy differences between training and task performance (Train accuracy – Task accuracy, plotted so that effects are primarily positive). **, significant after Bonferroni–Holm correction (p < 0.05).

Accuracy results followed a broadly consistent pattern. In the configural condition, accuracy significantly decreased from baseline across all tasks: inversion, t(19) = 27.23, p < .001; misalignment, t(19) = 17.51, p < .001; part–whole, t(19) = 9.85, p < .001; and composite, t(19) = 7.29, p < .001. For the featural condition, accuracy also declined in the inversion, t(19) = 8.17, p < .001, and misalignment, t(19) = 6.02, p < .001, tasks, though the magnitude of these effects was smaller than in the configural condition. In contrast, featural models showed reversed effects in the part–whole, t(19) = –2.46, p = .024, and composite, t(19) = –41.03, p < .001, tasks, indicating a clear opposite pattern relative to configural models. Between-condition comparisons again revealed reliable differences for inversion, t(19) = 9.30, p < .001; misalignment, t(19) = 9.00, p < .001; part–whole, t(19) = 9.28, p < .001; and composite, t(19) = 32.93, p < .001 (Fig. 4b). All significant results reported above remained robust after Bonferroni–Holm correction.

## 3. Experiment 2: 1^st^ order configuration

### 3.1 Human behavior

#### 3.1.1 Methods

##### 3.1.1.1 Participants

Forty-five undergraduate students from OSU participated in the inversion and misalignment tasks, and forty others participated in the part-whole and composite tasks. All participants were recruited through the REP and received course credit for their participation. Only those who achieved an accuracy rate of at least 80% in the last 30 trials of the preliminary training phase were included in the final analysis, resulting in a final sample of 20 participants for each set of tasks.

##### 3.1.1.2 Stimuli

The 1st-order configural and featural stimuli were created using the same specifications as in Section 2.1.1.2: all stimuli were generated using MATLAB and displayed on a 23.8-inch Dell monitor (1920×1080 resolution). Each stimulus featured a pentagonal frame containing five symbols (1167×875 pixels), including two fixed “●” placeholders to ensure consistent positional structure across conditions. The remaining three symbols determined group membership and were distinct across configural and featural conditions.

- **Configural Condition (Fig. 5a):** Each stimulus included the same three symbols (“∇”, “*”, “ω”), with group classification based solely on their spatial arrangement—i.e., 1st-order configural information. Six unique permutations were created by varying the positions of the symbols within the pentagon while holding their identities constant. To eliminate reliance on local cues, each symbol appeared equally often in every position across groups. Thus, classification could not be accomplished by detecting any individual symbol or fixed location; instead, participants had to learn the spatial relationships among at least **two** symbols to distinguish group membership.
- **Featural Condition (Fig. 5b):** Stimuli in this condition matched the configural condition in spatial layout but used variable symbol identities. Each stimulus displayed three symbols drawn from a set of four (“©”, “±”, “♡”, “≈”), with grouping determined entirely by feature combinations. The presence of **both** “©” and “±” signaled Group 1, while **both** “♡” and “≈” indicated Group 2. A third symbol, randomly drawn from the opposite group, was included as a distractor. Because no single symbol was diagnostic, successful classification required attention to specific feature pairings—again encouraging reliance on **two** pieces of information rather than any single cue.

##### 3.1.1.3 Design and Procedure

All aspects of the design and procedure were identical to those described in Section 2.1.1.3 (Fig. 6b), except that participants performed either the inversion and misalignment tasks, or the part-whole and composite tasks (in the experiments above, participants performed all four tasks).

**Figure 5.**
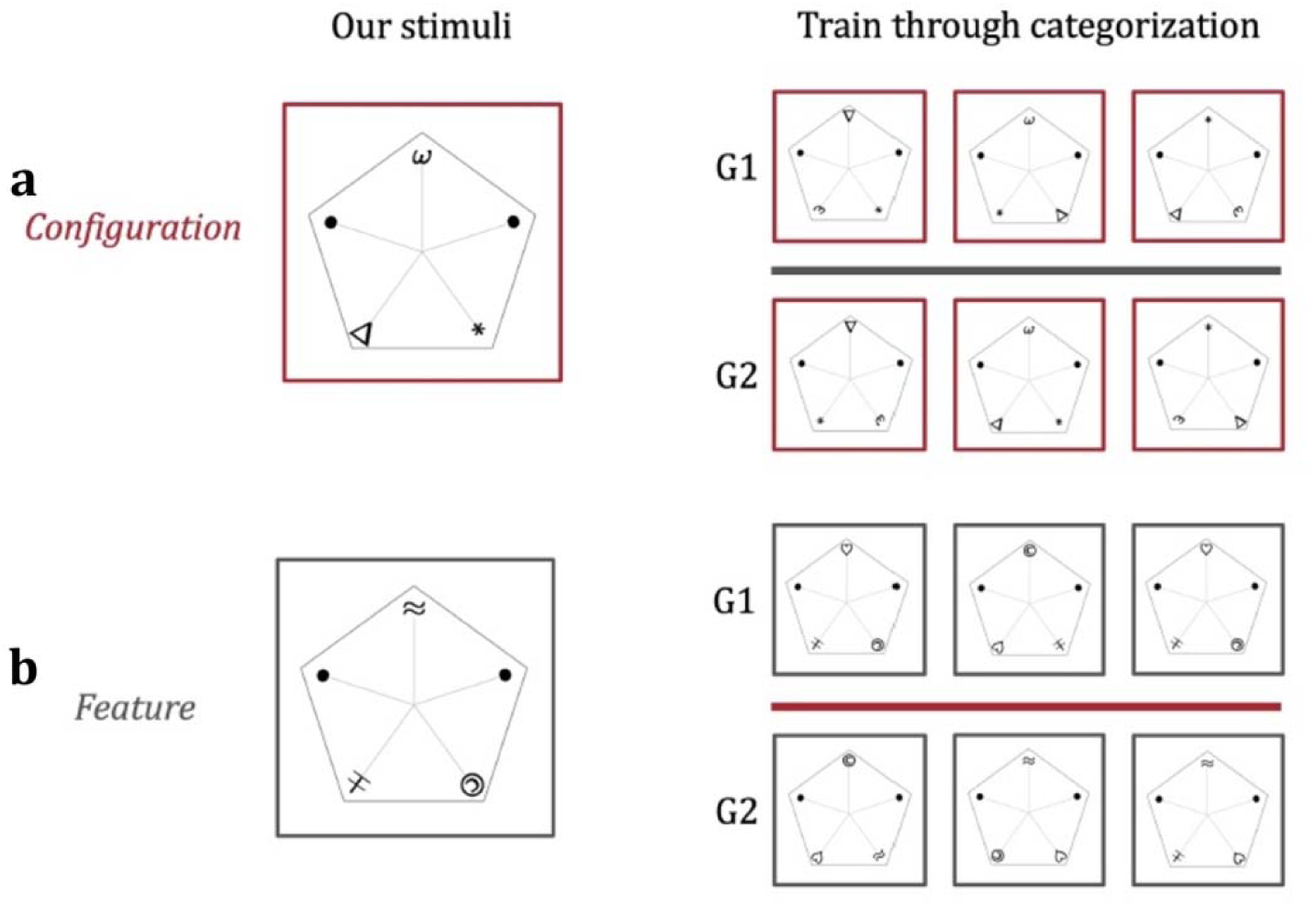
Examples of first-order stimuli (cropped into squares for better visualization). (a) Configural stimuli. Groupings were carefully designed so that the probability of each symbol appearing in any position was equal and consistent across groups. Thus, correct categorization required reliance on the relative configuration of at least two symbols. (b) Featural control stimuli. The grouping rule was identical to that of the second-order featural condition, depending on whether a specific pair of symbols co-occurred. Group 1 required the simultaneous presence of “©” and “±,” while Group 2 required the simultaneous presence of “♡” and “≈.” Since the third distractor symbol was drawn from the opposite group, correct categorization required recognition of at least two symbols.

**Figure 6.**
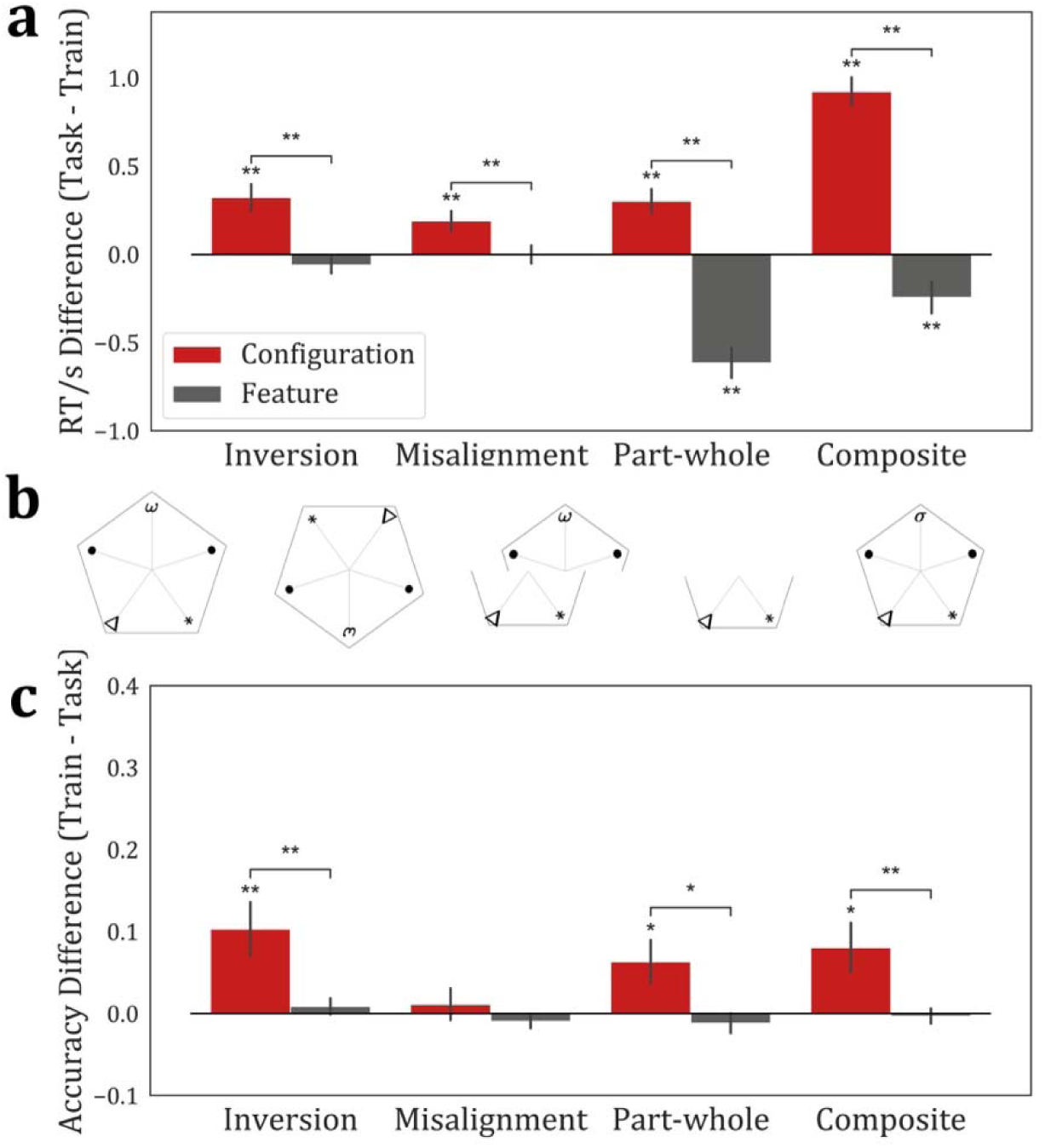
Behavioral results for first-order stimuli. (a) RT differences between training and task performance (Task RT – Train RT, plotted so that effects are primarily positive). (b) Example stimuli used during training and tasks. For the composite condition, one of the three symbols was randomly replaced with a novel symbol (illustrated here in the top position). (c) Accuracy differences between training and task performance (Train accuracy – Task accuracy, plotted so that effects are primarily positive). *, p < 0.05; **, significant after Bonferroni–Holm correction (p < 0.05).

#### 3.1.2 Results

At the end of the training phase (final 30 trials), no significant differences were observed between the configural and featural conditions in either RT or accuracy for both participant groups. Therefore, the configural and featural conditions are most likely of similar difficulty and the featural condition is an adequate control for the configural condition. For the inversion and misalignment group (n = 20), mean RTs were 1.25 s (SE = 0.08) for the configural condition and 1.29 s (SE = 0.09) for the featural condition, with no significant difference, t(19) = −0.62, p = .542. Accuracy was similarly high and comparable between conditions, with 96.00% (SE = 1.15%) for configural and 95.67% (SE = 0.97%) for featural, t(19) = 0.27, p = .789. For the part– whole and composite group (n = 20), mean RTs were 1.28 s (SE = 0.09) for configural and 1.35 s (SE = 0.11) for featural, t(19) = −0.68, p = .507. Accuracy was marginally higher in the featural condition (96.95%, SE = 1.10%) than in the configural condition (94.81%, SE = 1.10%), t(19) = −2.11, p = .048, although this does not survive corrections for multiple comparisons.

In the testing phase, participants showed clear holistic processing effects in the configural condition across all four tasks. RTs were significantly longer than the training baseline for inversion, *t*(19) = 4.27, *p* < .001; misalignment, *t*(19) = 3.30, *p* = .004; part–whole, *t*(19) = 4.44, *p* < .001; and composite, *t*(19) = 11.38, *p* < .001. None of these effects were not observed in the featural control condition. RTs in the featural condition did not differ significantly from baseline for inversion, *t*(19) = –1.28, *p* = .216, or misalignment, *t*(19) = –0.02, *p* = .985. Moreover, the part–whole and composite tasks in the featural condition showed significantly faster RTs compared to baseline, *t*(19) = –7.26, *p* < .001, and *t*(19) = –2.73, *p* = .013, respectively. Direct comparisons between the configural and featural conditions revealed significant RT differences in all tasks: inversion, *t*(19) = 4.47, *p* < .001; misalignment, *t*(19) = 3.21, *p* = .005; part–whole, *t*(19) = 10.46, *p* < .001; and composite, *t*(19) = 9.54, *p* < .001 (Fig. 6a).

Accuracy results showed a similar pattern. In the configural condition, accuracy in the testing phase was significantly lower than at baseline for inversion, *t*(19) = 3.14, *p* = .005; part–whole, *t*(19) = 2.40, *p* = .027; and composite, *t*(19) = 2.65, *p* = .016. However, the part–whole and composite effects did not survive Bonferroni–Holm correction, although they remained significant as independent tests. The misalignment effect was not significant, *t*(19) = 0.90, *p* = .380. In the featural condition, no accuracy differences from baseline were observed in any task: inversion, *t*(19) = 0.90, *p* = .380; misalignment, *t*(19) = –1.22, *p* = .238; part–whole, *t*(19) = –0.99, *p* = .333; and composite, *t*(19) = –0.36, *p* = .726. Pairwise comparisons between conditions revealed significant accuracy differences for inversion, *t*(19) = 2.97, *p* = .008, and composite, *t*(19) = 2.76, *p* = .013. The part–whole comparison, *t*(19) = 2.38, *p* = .028, did not survive Bonferroni–Holm correction, though it was significant as an independent test. The misalignment task showed no reliable difference between conditions, *t*(19) = 0.95, *p* = .356 (Fig. 6c).

### 3.2 Computer vision

#### 3.2.1 Methods

The methodology closely followed that of the previous CNN experiments (Section 2.2.1). The only difference was that the exemplars used to train and test the models were generated based on the new 1st-order configural and featural stimuli. As before, 5% random pixel noise was added to each exemplar to create a large set of augmented images for training and testing. The main analyses presented here were conducted using pre-trained ResNet-18 models. For results using other ResNet architectures, see Supplemental Figure S2.

#### 3.2.2 Results

In the model testing phase, classification probabilities in the configural condition significantly declined from baseline in all four tasks: inversion, *t*(19) = 15.30, *p* < .001; misalignment, *t*(19) = 22.46, *p* < .001; part–whole, *t*(19) = 10.34, *p* < .001; and composite, *t*(19) = 20.22, *p* < .001. In the featural condition, inversion and misalignment also produced significant changes, *t*(19) = 5.86, *p* < .001 and *t*(19) = 6.03, *p* < .001, respectively, though the effects were again smaller in magnitude than in configural networks. Both the part–whole, t(19) = −7.83, p < .001, and composite, t(19) = −10.22, p < .001, tasks showed significant reversed effects. Pairwise comparisons confirmed stronger effects in the configural condition for all tasks: inversion, *t*(19) = 6.16, *p* < .001; misalignment, *t*(19) = 11.35, *p* < .001; part–whole, *t*(19) = 16.26, *p* < .001; and composite, *t*(19) = 22.00, *p* < .001 (Fig. 7a).

**Figure 7.**
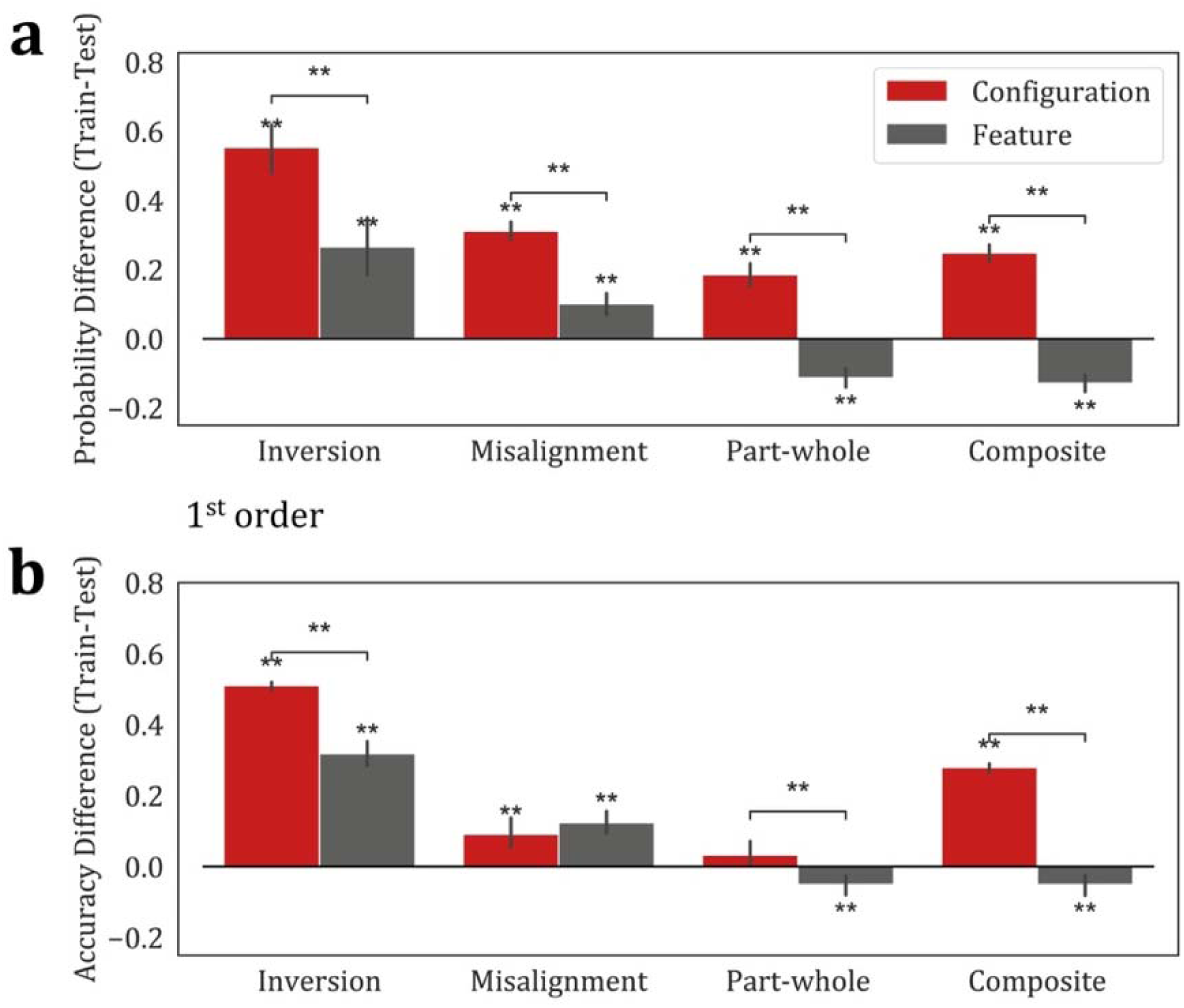
Computer vision results for first-order stimuli. (a) Probability differences between training and task performance (Train probability – Task probability, plotted so that effects are primarily positive). (b) Accuracy differences between training and task performance (Train accuracy – Task accuracy, plotted so that effects are primarily positive). **, significant after Bonferroni–Holm correction (p < 0.05).

Accuracy results showed a similar though somewhat weaker pattern. Configural models showed significant accuracy reductions in inversion, *t*(19) = 101.69, *p* < .001; misalignment, *t*(19) = 4.32, *p* < .001; and composite, *t*(19) = 47.11, *p* < .001. Accuracy in the part–whole task did not differ from baseline, *t*(19) = 2.00, *p* = .060. In featural models, inversion and misalignment again showed significant differences, *t*(19) = 17.47, *p* < .001 and *t*(19) = 7.34, *p* < .001, respectively, but reversed effect was found in both the part–whole, *t*(19) = −3.39, *p* = .003, and composite tasks, *t*(19) = −3.41, *p* = .003. Comparisons between conditions revealed significant differences in accuracy for inversion, *t*(19) = 9.95, *p* < .001; part–whole, *t*(19) = 3.62, *p* = .002; and composite, *t*(19) = 20.58, *p* < .001. No reliable difference was found in the misalignment task, *t*(19) = −1.19, *p* = .250 (Fig. 7b). All significant results reported above remained robust after Bonferroni–Holm correction.

## 4. Configuration and individual differences in face recognition

### 4.1 Methods

These experiments were performed online, and we recruited a total of 132 participants through REP and Prolific. Sixty-seven participants were excluded because their average accuracy on catch trials (see below) across tasks was below 50%, indicating that they were not properly attending or actively performing the online tasks. The final sample comprised 65 participants who were included in subsequent analyses.

For each participant, we collected data from the first- and second-order configuration training and task paradigms, as described in Sections 2.1.1.3 and 3.1.1.3, except that they also included a catch trial where the screen simply displayed, “Do not press any key”. In addition, we administered the Cambridge Face Memory Test (CFMT; Duchaine & Nakayama, 2006) to assess individual differences in face recognition ability.

### 4.2 Results

We examined correlations between CFMT accuracy and the strength of inversion, misalignment, part–whole, and composite effects, defined as the RT difference between task and training phases (Task RT – Train RT). Pearson’s correlations were computed for each effect.

For the first-order configural condition, none of the correlations reached significance, with *r* values close to zero (all *p* > .50). In the second-order configural condition, however, all four effects were positively and significantly correlated with CFMT performance: inversion (*r* = 0.37, *p* = .003), misalignment (*r* = 0.30, *p* = .017), part–whole (*r* = 0.37, *p* = .003), and composite (*r* = 0.32, *p* = .012). In the first-order featural condition, most effects were non-significant, although the misalignment effect showed a modest correlation (*r* = 0.26, *p* = .047), but did not remain significant after Bonferroni–Holm adjustment. The second-order featural condition likewise yielded no significant correlations (all *p* > .48). Full correlation results for all conditions are provided in Supplementary Table S1.

Taken together, significant associations with CFMT performance were observed only for the second-order configural effects across inversion, misalignment, part–whole, and composite tasks, whereas none of the first-order configural or featural effects remained significant after correction (Fig. 8).

**Figure 8.**
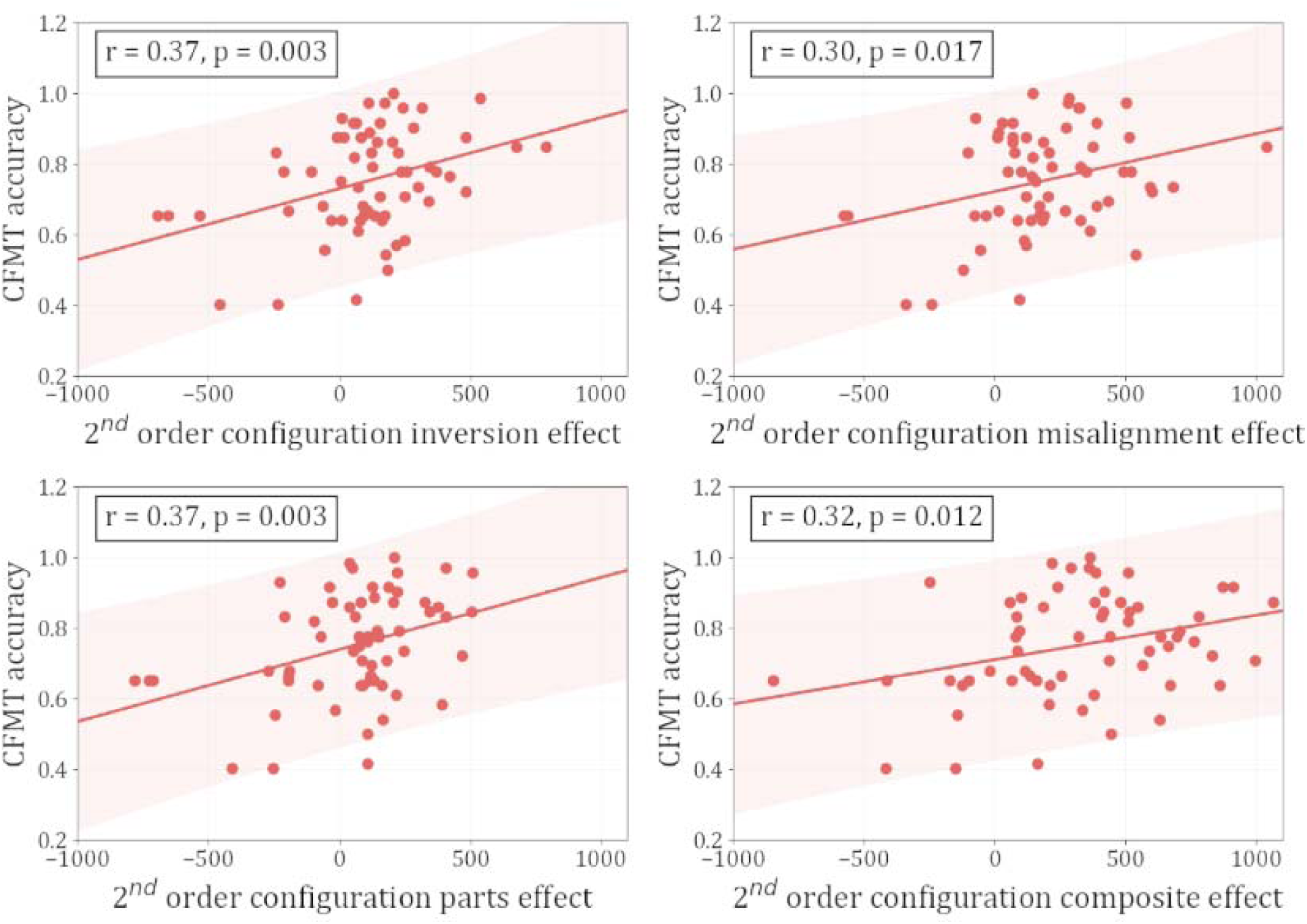
Correlations between second-order configural holistic effects and face-recognition ability (CFMT). Each panel shows CFMT accuracy (y-axis) as a function of the strength of the corresponding holistic effect (x-axis; Task RT – Train RT) for inversion, misalignment, part–whole, and composite. Points are individual participants (n = 65). The solid line is the ordinary least-squares fit, and the shaded band depicts the 95% prediction interval for observations. The boxed statistics report the Pearson correlation coefficient and the corresponding uncorrected p-value. All four second-order configural effects remain significant after Bonferroni–Holm correction..

## 5. Discussion

Our study set out to clarify two long-standing questions about holistic processing: (a) Is it truly unique to face perception? and (b) What type of information triggers it? Using carefully controlled, abstract novel stimuli, we provide converging behavioral and computational evidence that speak directly to both issues.

First, we asked whether holistic processing can arise outside the domain of faces. If holistic processing only arises for faces, then it might indeed reflect face-specific mechanisms. But if it can also occur for non-face stimuli, then we must ask: what properties of faces give rise to holistic processing in the first place? Perhaps the critical factor isn’t the “face-ness” of the stimulus, but rather something more general, such as the presence of certain types of spatial configuration. Using novel stimuli composed of symbolic elements and lacking any face-like structure, we observed robust inversion, misalignment, part–whole, and composite effects, each of which is a hallmark of holistic face perception. Critically, these effects emerged only when categorization relied on spatial relationships among features (i.e. configuration), not on feature identity alone. Our stimuli were deliberately designed to avoid perceptual face-likeness: we reversed the “top-heavy” property that characterizes faces and face-like patterns, a cue known to elicit face preferences even in newborns (Morton & Johnson, 1991; Valenza et al., 1996; Viola Macchi et al., 2004). They also lack curvature, color, and other mid-level visual features that are indicative of animacy (Long et al., 2017; Rosenthal et al., 2018; Schmidt et al., 2017). This design ensures that the observed holistic effects cannot be attributed to inadvertent face resemblance. In doing so, our results extend beyond prior work using face-like stimuli such as Greebles (Gauthier & Tarr, 1997), which have been noted for triggering face-specific processing due to their structural similarity to real faces (Brants et al., 2011). Taken together, our findings provide strong evidence that holistic processing is not exclusive to face perception and does not require activation of face-specific mechanisms (Farah et al., 1998; Maurer et al., 2002; McKone et al., 2007).

Second, we examined what triggers holistic processing in the first place. While traditional accounts have emphasized second-order configural information, defined as the precise spatial distances between features, as essential, our findings challenge this assumption. We observed equally strong holistic effects when only first-order spatial relationships were available (i.e., categorical information about which symbol was above, below, or beside another). As long as categorization required integrating at least two spatial relations, holistic processing emerged. In contrast, when classification could be achieved based solely on the presence of specific feature combinations, no holistic effects were found. These results identify spatial configuration itself, rather than feature content or long-term expertise, as the primary trigger for holistic processing.

Importantly, our participants acquired the necessary configural representations in a brief training period (∼30 minutes), far shorter than the extended experience typically required to induce holistic effects in non-face domains (Diamond & Carey, 1986; Gauthier & Tarr, 1997). This suggests that expertise is not a necessary condition. Rather, once participants have learned to rely on spatial relations for categorization, holistic effects readily emerge. That said, our findings do not rule out the possibility that expertise may amplify or refine holistic processing; it simply is not required. Future work could test whether prolonged training in a featural task, in which configuration is irrelevant, could eventually lead to holistic processing once spatial relationships are internalized.

Notably, we observed a similar pattern in convolutional neural networks. CNNs trained on the configural condition displayed the full suite of holistic effects, closely mirroring those seen in humans, whereas CNNs trained on the featural condition did not. Previous work with CNNs has focused largely on holistic responses to faces (Dobs et al., 2023; Peng et al., 2021; Tian et al., 2022; Wirth et al., 2024). Here, we demonstrate for the first time that CNNs can develop holistic processing for non-face stimuli when trained to rely on configuration. This parallel across artificial and biological systems suggests that holistic processing may arise naturally whenever learning is driven by spatial configuration. As such, CNNs offer not only functional analogues of human perception but may also serve as reverse-engineering tools to probe the computational foundations of holistic vision (Huang et al., 2022; Kanwisher et al., 2023; Lu & Ku, 2023).

While our findings challenge the idea that holistic processing is face-specific, they do not imply that face perception is not a special visual category, nor that the prior reports of holistic processing of faces are incorrect. Faces continue to evoke distinctive neural signatures, such as the N170 ERP component (Bentin et al., 1996) and consistent activity in the fusiform face area (Kanwisher et al., 1997). From a behavioral perspective, faces may simply be the most configuration-rich category we routinely encounter. Because faces require precise relational judgments for identity recognition, they reliably elicit holistic processing (Rhodes et al., 1993; Robbins & McKone, 2007).

Further, we examined whether holistic processing is related to face-recognition ability. Previous research has demonstrated a relationship between these effects and face recognition ability: Wang et al. (2012) found that part–whole and composite effects predicted face recognition, and Rezlescu et al. (2017) argued that the inversion effect was a reliable predictor. In the present study, by using non-face stimuli to probe configural processing, we eliminated the confound of face information appearing in both the predictor and outcome measures. Strikingly, we obtained a consistent pattern: all four classic paradigms of holistic processing predicted face-recognition ability, but only when they involved second-order configuration, and feature-based recognition does not produce these effects. This finding aligns with the view that second-order configural processing is the component of holistic processing most closely tied to face recognition.

Our findings also point toward several directions for future research. One limitation is that participants learned to categorize stimuli into two groups, rather than individuate specific exemplars. Prior studies suggest that holistic processing is often enhanced by individuation demands, particularly in faces and expert object categories (Kanwisher, 2000; Wong et al., 2009). Investigating whether individuation modulates holistic effects in our stimuli could further clarify the relationship between task demands and processing style. In addition, we did not collect eye-tracking data. Given that holistic perception may involve distinctive gaze patterns or feature-integration strategies, eye-tracking could offer a richer picture of how observers allocate attention when processing configural versus featural information (Kundel et al., 2007; Zhong et al., 2023).

Lastly, our findings have potential clinical relevance. Deficits in holistic processing have been implicated in prosopagnosia and autism spectrum disorders (Behrmann et al., 2006; Palermo et al., 2011). By isolating configural and featural information in a controlled, domain-general format, our stimuli may provide a useful diagnostic tool for probing these impairments and for studying their neural correlates. Recent computational research also suggests that enhancing configural structure improves object recognition in artificial systems (Jang et al., 2024), reinforcing the notion that configural processing is a foundational component of visual cognition, in both natural and artificial.

In sum, our study demonstrates that holistic processing is not a privilege of faces, but a general response to stimuli structured around spatial configuration. Whether the information is fine-grained or categorical, human and artificial systems alike exhibit holistic processing once they are trained to rely on spatial relationships. This highlights the need to better understand configuration as a central feature of visual processing and cautions against treating holistic processing as a signature exclusive to face perception.

## Supplemental

**Figure S1.**
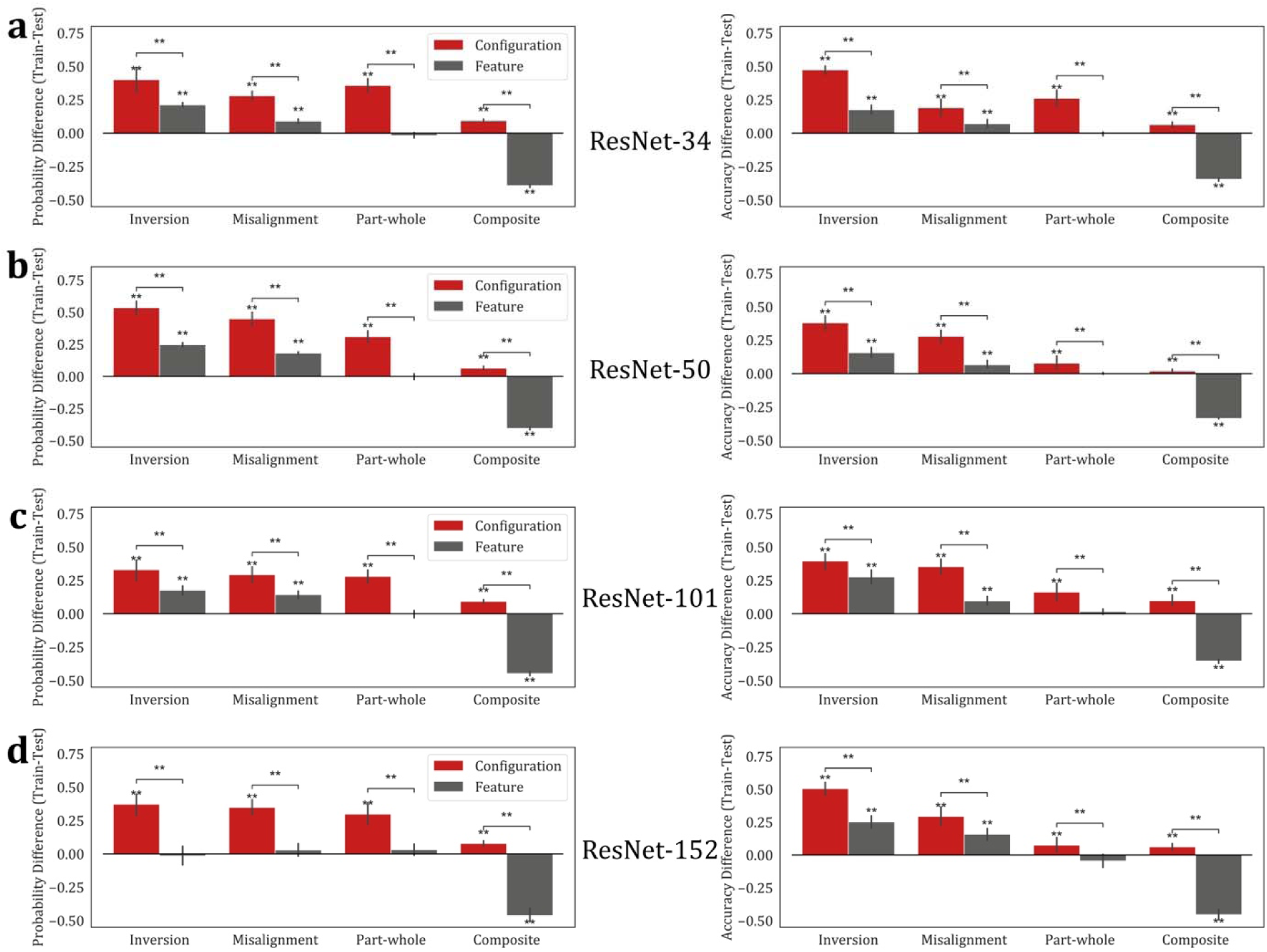
Computational results for second-order stimuli across different ResNet architectures. Left panels show probability differences between training and task performance (Train probability – Task probability, plotted so that effects are primarily positive). Right panels show accuracy differences between training and task performance (Train accuracy – Task accuracy, plotted so that effects are primarily positive). Results are presented for ResNet-34 (a), ResNet-50 (b), ResNet-101 (c), and ResNet-152 (d). **, significant after Bonferroni–Holm correction (*p* < 0.05).

**Figure S2.**
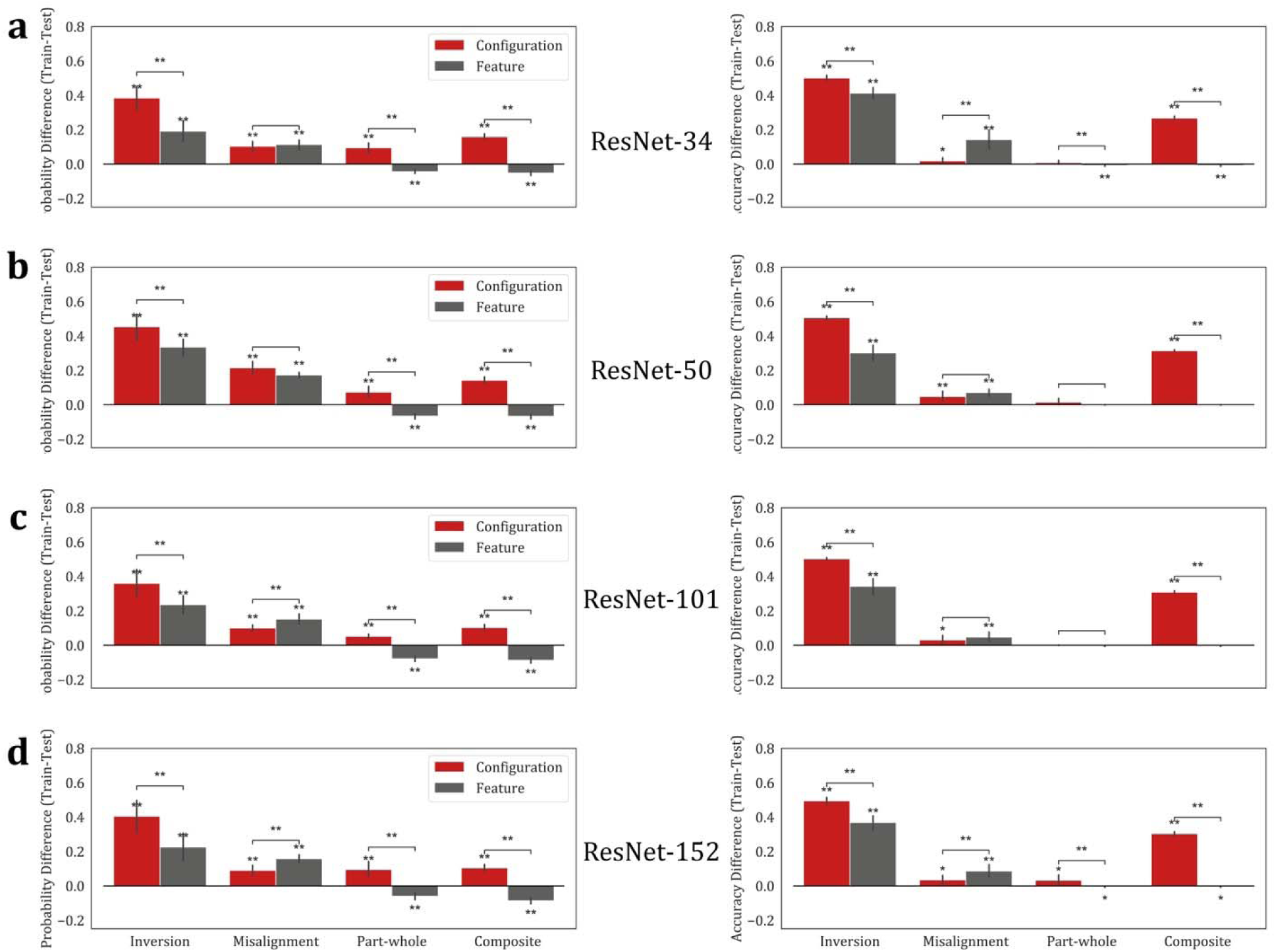
Computational results for first-order stimuli across different ResNet architectures. Left panels show probability differences between training and task performance (Train probability – Task probability, plotted so that effects are primarily positive). Right panels show accuracy differences between training and task performance (Train accuracy – Task accuracy, plotted so that effects are primarily positive). Results are presented for ResNet-34 (a), ResNet-50 (b), ResNet-101 (c), and ResNet-152 (d). *, *p* < 0.05; **, significant after Bonferroni–Holm correction (*p* < 0.05).

---

**Table S1.** Correlations between CFMT accuracy and the strength of holistic effects (Task RT – Train RT) across conditions. Shown are Pearson’s correlation coefficients and uncorrected *p*-values for inversion, misalignment, part–whole, and composite effects in first- and second-order configural and featural conditions. * *p* < .05; ** significant after Bonferroni–Holm correction (*p* < .05).

| *Effect* | *2ndOrder*<br>*Configuration* | *2ndOrder*<br>*Feature* | *1stOrder*<br>*Configuration* | *1stOrder*<br>*Feature* |
| --- | --- | --- | --- | --- |
| *Inversion* | r=0.368<br>p=0.003** | r=0.090<br>p=0.491 | r=-0.017<br>p=0.898 | r=-0.034<br>p=0.798 |
| *Misalignment* | r=0.302<br>p=0.017** | r=-0.043<br>p=0.743 | r=-0.057<br>p=0.668 | r=0.260<br>p=0.047* |
| *Part-Whole* | r=0.371<br>p=0.003** | r=0.091<br>p=0.487 | r=0.011<br>p=0.934 | r=0.063<br>p=0.637 |
| *Composite* | r=0.316<br>p=0.012** | r=-0.011<br>p=0.934 | r=0.083<br>p=0.533 | r=0.094<br>p=0.478 |

